# Diesel exhaust particles induce lasting and age-dependent damage to the brain in *Drosophila melanogaster*

**DOI:** 10.1101/2025.08.20.671202

**Authors:** Ranchana Yeewa, Siwat Poompouang, Kornravee Photichai, Titaree Yamsri, Natsinee U-on, Wasinee Wongkummool, Phatsara Manussabhorn, Luca Lo Piccolo, Salinee Jantrapirom

## Abstract

Diesel exhaust particles (DEP), major air pollutants emitted from automobile engines, contain numerous toxic compounds. While the adverse effects of DEP exposure on the respiratory and cardiovascular systems are well documented, its impact on brain health remains poorly understood. In this study, we employed Drosophila melanogaster as a model organism to investigate the neurological effects of DEP exposure and the impact of exposure cessation across different age groups. Molecular, histopathological, and behavioral markers were assessed before and after exposure to varying doses of DEP at different time intervals.Interestingly, DEP exposure induced age-dependent cellular responses in the brain, including elevated reactive oxygen species (ROS), increased neuroinflammation, and disruption of the blood-brain barrier (BBB). Prolonged exposure led to pronounced vacuolization in the brains of aged flies. While cessation of DEP exposure resulted in partial recovery in young flies particularly when implemented early, aged flies exhibited limited benefit, with persistent evidence of likely irreversible brain damage. Overall, this study invites greater public awareness and careful consideration in public health policy to limit long-term DEP exposure, particularly among older individuals, and to encourage strategies that reduce potential risks to brain health associated with air pollution.

**Highlights:**

1) DEP exposure is detrimental to the brain.
2) The brain responds to DEP exposure in an age-specific manner.
3) Permanent damage to the brain of old flies results from DEP exposure.
4) Cessation mitigates DEP-induced brain impairments when implemented at a young age.

**Environmental Implication:** Our study highlights the detrimental, age-dependent effects of diesel exhaust particle (DEP) exposure on brain health, underscoring the urgency of reducing air pollution. The findings support stricter environmental regulations to limit DEP emissions and promote cleaner transportation alternatives. Protecting vulnerable populations, particularly the elderly, from prolonged exposure may help mitigate the long-term neurological impacts of air pollutants and reduce the public health burden.

## Introduction

The significant health risks posed by air pollution underscore the urgent need for enhanced public health measures to mitigate its adverse effects. Studies conducted by the International Council on Clean Transportation (ICCT) estimate that approximately 385,000 premature deaths globally are attributable to air pollution from vehicle exhaust emissions, particularly those from diesel engines (1). This statistic highlights the profound global health impact of diesel exhaust emissions, necessitating stringent regulatory actions. Many countries have implemented measures to limit diesel exhaust particle (DEP) concentrations and exposure durations, particularly in occupational settings where direct exposure is common (2, 3). Nevertheless, exposure to DEP remains a significant concern, particularly in urban centers with high traffic congestion (4).

DEP represents a complex mixture of gaseous and particulate components, consisting of ultrafine carbon particles coated with carcinogenic compounds and metals (5–7). Extensive research has linked DEP exposure to elevated risks of respiratory and cardiovascular diseases, various cancers, and premature mortality (8–13). Among these, the neurological effects of DEP have garnered increasing attention due to the limited understanding of causative mechanisms and the specific pathways involved.

Controlled human exposure studies have shown that acute exposure to DEP induces alterations in brain activity, notably an increase in β2 fast-wave activity in the frontal cortex. This pattern is associated with cortical stress and is commonly observed in neurological and neuropsychological disorders (14, 15). Additionally, prolonged DEP exposure has been linked to cognitive impairments, particularly in children and adolescents. While neuroinflammation and oxidative stress have been proposed as key mediators of these neurological effects, further research is required to elucidate deeply the underlying mechanisms (16–18). Moreover, gaps remain regarding the effects of DEP exposure in both young and aging populations. Specifically, the differential responses of these age groups to short– and long-term exposure cessation, along with the associated risks and potential recovery benefits, are poorly understood. Given the logistical and ethical challenges of conducting long-term human studies, animal models offer a valuable alternative for addressing these questions.

Toxicological studies have long complemented epidemiological research by providing biological plausibility for the health effects of DEP. Traditional mammalian *in vivo* models, which closely replicate human physiology, have been instrumental in studying the impacts of inhalation, the primary route of exposure (19–21). However, these models face limitations, particularly in high-throughput screening and the investigation of long-term effects, especially in aging populations. To address these challenges, alternative model organisms such as zebrafish, nematodes, and *Drosophila melanogaster* have emerged as valuable tools. These models offer the advantage of being fully intact *in vivo* systems while providing the logistical simplicity, scalability, and efficiency typically associated with *in vitro* assays.

Here, we leverage the utility of *Drosophila melanogaster* to investigate the neurological impacts of DEPs exposure across the lifespan, focusing on young and aging populations. This study examines the effects of DEP on the nervous system and explores the potential recovery and benefits following exposure cessation in these distinct age groups.

## Materials and methods

### Drosophila husbandry

The wild-type *Oregon R* strain (*Oregon R*) (BDSC #2376, USA) was obtained from the Bloomington *Drosophila* Stock Center (USA). Flies were maintained on standard cornmeal-yeast-glucose medium under controlled conditions of 25°C, 55% humidity, and a 12:12 light/dark cycle, unless specified otherwise.

### Diesel Exhaust Particle (DEP) Exposure

Standard diesel exhaust particles (DEPs; Industrial Forklift, NIST2975) were obtained from the National Institute of Standards and Technology. A DEP stock solution was prepared by dissolving the particles in 100% dimethyl sulfoxide (DMSO) (CAS No. 67-68-5; RCI Labscan, Thailand) at a concentration of 20 mg/mL. The solution was sonicated for at least 20 minutes to ensure thorough dispersion before being mixed into the fly food medium. For experimental use, the stock was diluted into standard fly food to achieve final concentrations of 1, 10, 100, or 200 μg/mL (hereafter referred to as DEP1, DEP10, DEP100, and DEP200, respectively). Young adult flies (three-day-old males; D3) were maintained on DEP-containing food for the duration of the experiments. Aged flies were first maintained on standard food until 30 days of age, after which they were transferred to food containing the assigned DEP concentrations. In all experiments, flies were transferred to fresh food every 3–4 days until the end of the experimental period. For larval experiments, parental crosses were set up in vials containing the appropriate DEP concentration, and the resulting embryos were raised until the third instar foraging larval stage (L3). Three exposure durations were used in this study, namely acute, prolonged and chronic corresponding to 7, 14 and longer than 15 days. Unless otherwise stated chronic exposure was provided troughtout the life span.

### Gene expression analysis

RNA isolation and quantitative real-time PCR (qRT-PCR) were performed following previously established methods (22) with slight modifications. Briefly, total RNA was extracted from 40 adult fly heads per treatment group using the RNeasy® Mini Kit (Cat. No. #74106 QIAGEN, Germany). RNA purity and concentration were assessed by NanoDrop Spectrophotometer (ThermoFisher, USA) at 260/230 and 260/280 ratios, and 0.6 μg of total RNA was subsequently reverse-transcribed into cDNA using the SensiFAST cDNA Synthesis Kit (Cat. No. #BIO-65054, Bioline, United Kingdom). Gene expression changes were assessed using qRT-PCR performed in triplicate for each RNA extraction, with reactions prepared using SensiFAST SYBR® Lo-ROX Kit (Cat. No. #BIO-94005, Bioline, United Kingdom) and specific primer sets (Supplementary Table S1) on a CFX Opus 96 Real-Time PCR System. Relative mRNA levels were normalized to *ACT5C* expression using the comparative Ct method.

### Larval crawling assay

The crawling assay was conducted to assess larval locomotion, following the standardized protocol (23). Larval movement and crawling trajectories were monitored and analyzed using ImageJ version 1.53 K, integrated with the wrMTrck plugin, which facilitates precise tracking of movement patterns. The distance traveled by larvae within a 30-second interval was quantified and visualized the data for comparative analysis. The experiment was performed in at least three independent replicates, analyzing a minimum of 30 larvae per condition.

### Adult negative geotaxis

A climbing assay, recognized as a standard method for assessing the locomotive function of adult flies, was conducted following established protocols (24) of 10–12 adult male flies from each treatment group were placed in glass tubes (15 cm in height, 2.2 cm in opening width) without anesthesia to preserve natural behavior. The flies were tapped to the bottom of the tubes and given 30 seconds to climb vertically; this process was repeated five times and recorded on video for subsequent analysis. Individual climbing heights were scored within a 5-second interval using a predefined scale: 0 (less than 2.0 cm), 1 (2.0–3.9 cm), 2 (4.0–5.9 cm), 3 (6.0–7.9 cm), 4 (8.0–9.9 cm), and 5 (greater than 10 cm). The climbing index, quantifying overall locomotor performance, was calculated by multiplying the number of flies in each score category by their respective scores, summing the products, and dividing by the total number of flies in the group. The experiment was performed in at least three independent replicates, analyzing a minimum of 100 flies per condition.

### Lifespan assay

The viability of male flies was evaluated using the established method described in (22–24). All treatment and vehicle control groups were transferred to new vials containing the fresh medium every three to four days without using anesthesia to minimize the risk of acute mortality, particularly in aged flies. The number of dead flies in each group was recorded during each vial transfer to monitor survival trends. Flies that escaped or died from accidental causes were categorized as right-censored (not recorded as dead), as their deaths were unrelated to the treatment conditions, ensuring the accuracy of survival analysis. This process was repeated at regular intervals until all flies in each group had reached the end of their lifespan. The survival probability for each group was visualized using a Kaplan-Meier survival curve, plotting the percentage of surviving flies against the number of days elapsed. The experiment was conducted in five independent replicates, each group comprising at least 200 flies.

### *In vivo* detection of O₂•⁻ radical formation

The protocol for *in vivo* measurement of O₂•⁻ radical formation using dihydroethidium (DHE) staining was adapted from the previous study with modifications (25). Briefly, adult fly brains were dissected in ice-cold Schneider’s insect medium (Cat No. P04-91500, PAN Biotech, Germany). The dissected brain tissues were then incubated in 1× PBS containing 30 µM DHE at 25°C for 5 minutes under dark conditions. Following the staining procedure, the samples were fixed in 4% paraformaldehyde (PFA) at 25°C for 5 minutes, washed three times with 1× PBS, and mounted onto glass slides with spacers using Vectashield mounting medium. Samples were visualized using an Olympus FV3000 Confocal Laser Scanning microscope (Olympus, Japan) within 10 minutes of preparation. The intensity of DHE staining was quantified using Olympus cellSens Imaging Software.

### Blood-brain barrier (BBB) Injection Assay

The protocol was adapted from Axelrod, Li et al. 2023 (26), with modifications. Briefly, CO₂-anesthetized adult flies were microinjected using a FemtoJet 4i microinjector coupled with an InjectMan 4 micromanipulator (Eppendorf, Germany). Femtotip II capillaries (0.5 µm inner diameter, 0.7 µm outer diameter; Eppendorf, Germany) containing Dextran-conjugated Tetramethylrhodamine (TMR) dyes (molecular weight 10,000 kDa; D1868, Thermo Fisher, USA) were used for injections under a light stereomicroscope with controlled pressure and injection time parameters (injection pressure [Pi]: 180 hPa, compensation pressure [Pc]: 50 hPa, injection duration [ti]: 1.0 s). The dye was diluted in injection buffer (1X phosphate-buffered saline, 2.5 mM final concentration). For BBB permeability assessments, the dyes were injected into the lateral thorax region between the wing socket and haltere. Following injection, flies were allowed to recover for 30 minutes in dark chamber before decapitation and imaging using an Olympus FV3000 Confocal Laser Scanning microscope (Olympus, Japan). Laser and acquisition settings were standardized across all samples within the same experiment. Confocal stacks comprising 15–22 slices (10 μm) were acquired and processed into maximum projection images. Average pixel intensity in the eye region was quantified using Olympus cellSens Imaging Software. Experimental data were normalized to mock-injected controls. All injections were performed during the Zeitgeber time (ZT) 0.5–1.5 unless otherwise specified.

### Histological analysis of the brains

The dissection of flies’ heads and subsequent tissue processing were conducted according to the established protocol (27), which was recognized for its effectiveness in preserving adult brain morphology. Following overnight decalcification in 100% Formic acid (Cas No. #64-18-6, RCI Labscan, Thailand), the flies’ heads were carefully transferred to a mesh bag, and placed into a tissue processing cassette. The cassettes were then positioned in beakers to initiate manual tissue processing and embedding, ensuring careful and precise handling throughout the procedure. The embedded tissue was sectioned at 3.5 µm using a Microm HM 325 Microtome (Thermo Fisher, USA) and stained with Hematoxylin and Eosin (H&E). Next, the stained brain section images were acquired using a high-resolution Olympus BX53 Microscope with a digital imaging system. Brain degeneration, characterized by vacuolation, was identified as purple staining in the cell body region or pink staining in the neuropil region under H&E staining (Kretzschmar, Hasan et al. 1997). To ensure statistical reliability and consistency for the quantification of vacuolar neurodegeneration, the number of vacuoles throughout the entire brain was counted per fly and the average number of vacuoles per brain was averaged for each group (n ≥ 6) (Torre, Bukhari et al. 2023), providing a robust dataset for assessing neuronal and neuropil deterioration.

### Data analysis

Statistical analyses were performed using GraphPad Prism 9 (GraphPad Software). Comparisons between two groups were conducted using the Mann–Whitney U test, while comparisons among multiple groups were analyzed using the Kruskal–Wallis test followed by Dunnett’s post hoc test. For lifespan assays, survival curves were generated using the Kaplan–Meier method and compared using the log-rank (Mantel–Cox) test. Statistical significance was set at *p < 0.05*. All data, except for the lifespan analysis, are presented as mean ± standard deviation (SD).

## Results

### Acute DEP exposure triggers a cellular response primarily in the brain

Inflammation and oxidative stress are key mechanisms underlying the toxic effects of airborne pollutants (28, 29). To gain deeper insights into both systemic and neural responses, we assessed the impact of acute (7-days) exposure to various concentrations of DEP in young flies, focusing on both brain-specific and whole-body responses **(Figure 1A).**

**Figure 1.**
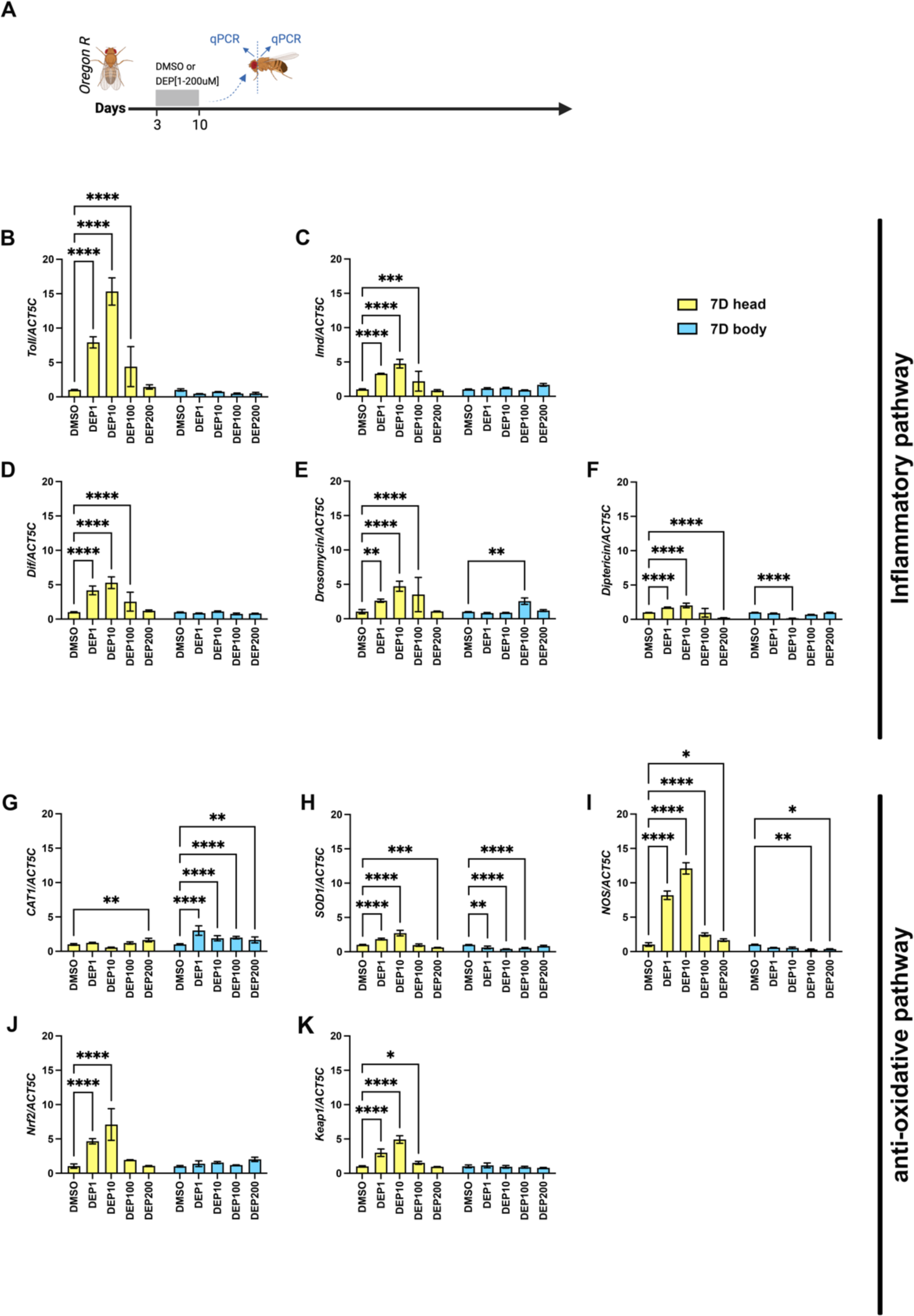
Effects of diesel exhaust particles (DEPs) on inflammatory and antioxidant responses in the heads and bodies of young flies following acute exposure. (A) Schematic of acute DEP exposure (7 days) and analysis timeline. (B–K) Relative mRNA expression of inflammatory markers including *Toll* (B), *Imd* (C), *Dif* (D), *Diptericin* (E) and antioxidant genes *CAT* (G), *SOD1* (H), *NOS* (I), *Nrf2* (J), *Keap1* (K) in the heads (yellow bars) and bodies (blue bars) of flies exposed to DEP at 1, 10, 100, or 200 µg/mL (DEP1–DEP200), compared to vehicle control (DMSO). Gene expression was normalized to *ACT5C*. Each condition included a minimum of three biological replicates. Statistical significance was assessed using the Kruskal–Wallis test with Dunn’s multiple comparisons. *\*p < 0.05, **p < 0.01, ***p < 0.001, ****p < 0.0001*.

In the brain, inflammatory markers such as *Toll* and *Imd*, along with their downstream targets, showed a dose-dependent increase beginning at 1 µg/mL (DEP1) and peaking at 10 µg/mL (DEP10), with no further increase at higher concentrations **(Figure 1B–C).** Antioxidant response genes—including *SOD1*, *NOS*, *Nrf2*, and *Keap1*—exhibited a similar pattern **(Figure 1H–K).**

In contrast, the whole-body response differed markedly: *mRNA* expression of inflammatory and antioxidative markers showed no consistent dose-dependent pattern, and most changes were not significantly different from vehicle-treated controls **(Figure 1B–K)**. Importantly, the different vehicle concentrations used in the food were also tested and showed no significant effect on inflammatory or oxidative pathways **(Supplementary Figure S1A–K).** These findings indicate that, under our experimental conditions, DEP exposure exerts a more pronounced effect on the brain than the rest of the body. This validates our experimental design as a reliable model for studying brain-specific effects of DEP exposure while minimizing systemic confounding. Based on these observations, we focused our subsequent experiments on the brain-specific effects of DEP at 10 and 100 µg/mL concentrations (DEP10 and DEP100, respectively).

### Acute DEP exposure increases ROS levels in the brain and affects locomotion in young flies

The brain is a metabolically active organ where reactive oxygen species (ROS) are continuously generated as byproducts of cellular metabolism (30). Under physiological conditions, antioxidative defenses maintain redox homeostasis by counterbalancing ROS production (31). However, excessive ROS can lead to oxidative stress, which is associated with neuronal damage (32).

Given that ROS not only mediate oxidative damage but also act as upstream regulators of inflammation and antioxidant responses, we next investigated whether DEP exposure directly induces oxidative stress in the brain. Accordingly, superoxide levels in the brains of flies following acute DEP exposure were measured using dihydroethidium (DHE) staining **(Figure 2A).** A significant increase in ROS levels was observed in flies exposed to DEP10, with a much more pronounced elevation in those exposed to DEP100 **(Figure 1B–F).**

**Figure 2.**
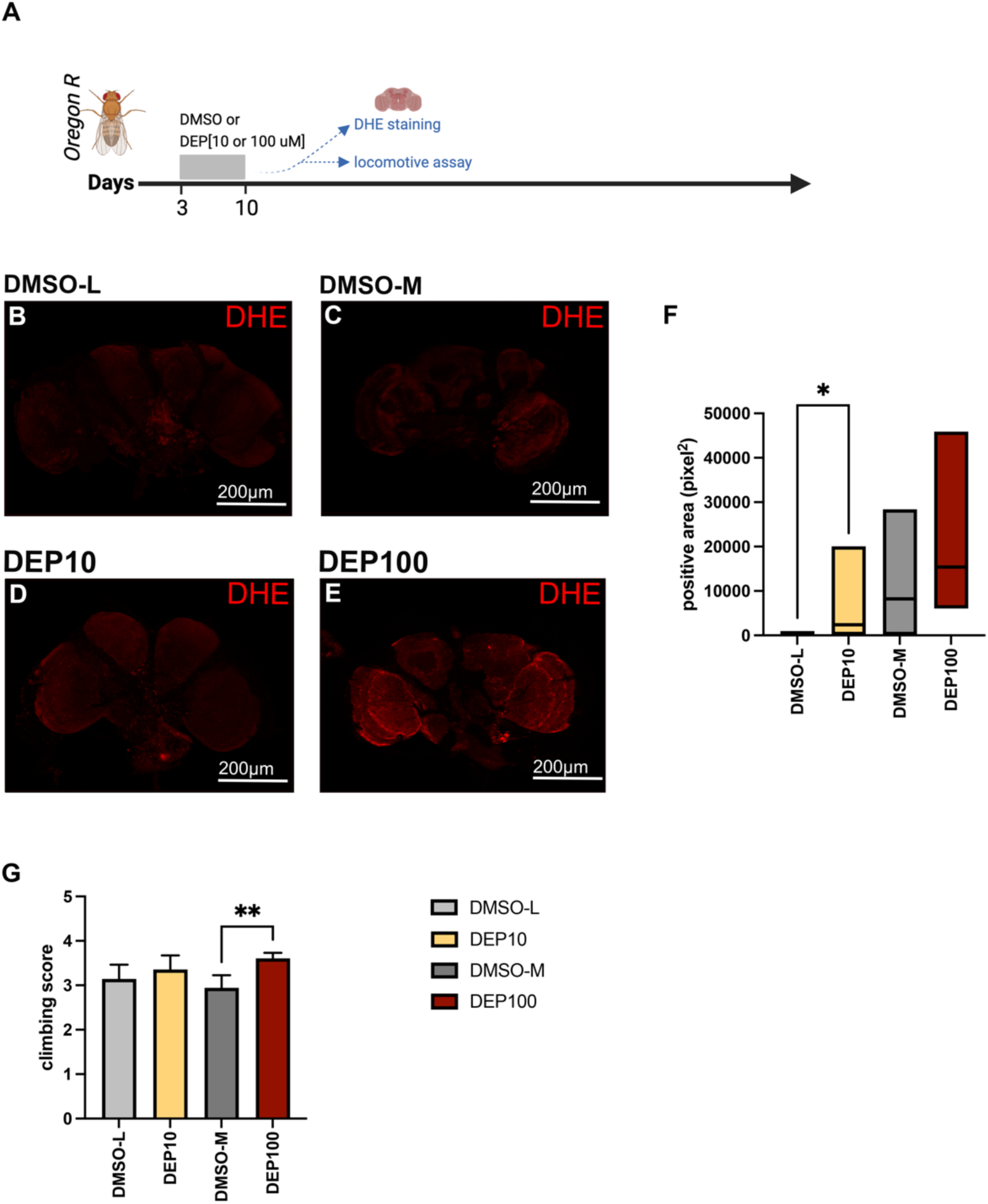
Effects of Diesel Exhaust Particles (DEPs) on young flies following acute exposure. (A) Schematic of acute DEP exposure (7 days) and analysis timeline. (B-E) Representative images showing dihydroethidium (DHE) staining in fly brains after acute exposure to (B) DMSO-L, (C) DMSO-M (D) DEP10, and (E) DEP100, indicating oxidative stress levels. (F) Quantification of DHE-positive area in each condition. Locomotor performance following DEP exposure (M) Adult locomotion after 7 consecutive days of DEP exposure. Each experiment included a minimum of three biological replicates. Statistical significance was assessed using the Kruskal–Wallis test with Dunn’s multiple comparisons. A p-value of < 0.05 was considered statistically significant *(*p < 0.05, **p < 0.01*).

As elevated ROS levels have been associated with altered locomotor function in flies (33, 34), we next evaluated climbing ability following acute DEP treatment. Exposure to either low or medium concentrations of DMSO (DMSO-L and DMSO-M) did not affect locomotor performance in young flies over a 14-day monitoring period (prolonged exposure) **(Supplementary Figure S2A)**. Similarly, DEP10 was well tolerated, with no significant changes in climbing ability **(Supplementary Figure S2D).** In contrast, exposure to DEP100 resulted in a noticeable increase in locomotor activity at the initial time point **(Supplementary Figure S2E)**. Notably, this hyperactivity was also observed when compared to both age– and vehicle-matched controls **(Figure 2G).**

To determine whether a similar behavioral response occurs at a different developmental stage, we conducted a crawling assay using third instar larvae. Larvae were exposed to DEP10, DEP100, or their respective vehicle controls, and motor function was assessed through crawling distance **(Supplementary Figure S3A)**. Under these experimental conditions, acute exposure to DEP100 also had a pronounced effect on locomotor function, with larvae covering significantly greater distances compared to vehicle-matched controls **(Supplementary Figure S3B).**

Altogether, short-term exposure to DEP100 alters locomotor behavior across developmental stages, and this behavioral change is accompanied by elevated ROS levels in the brain.

### Brain responds to acute DEP exposure in an age-specific manner

Several lines of evidence indicate that environmental pollutants can accelerate aged-related cellular decline and contribute to age-related diseases (35, 36). These findings prompted us to investigate the effects of DEP exposure on aged flies. Flies were raised on chemical-free food until they reached 30 days of age (D30), considered old in *Drosophila*. At this point, D30 flies were divided into four groups: those exposed acutely to DEP10 or DEP100, and those receiving only the vehicle (DMSO-L or DMSO-M). After exposure, we collected brains for RNA extraction and ROS measurement **(Figure 3A).** In contrast to young flies, markers of neuroinflammation and antioxidative genes in aged flies did not show significant changes in response to DEP exposure, with only *Toll* and *NOS* being statistically elevated in the heads of DEP10-treated flies **(Figure 3B–C)**. As aging itself can influence the expression of these genes, the additional effects triggered by DEP exposure may be less pronounced in aged flies.

**Figure 3.**
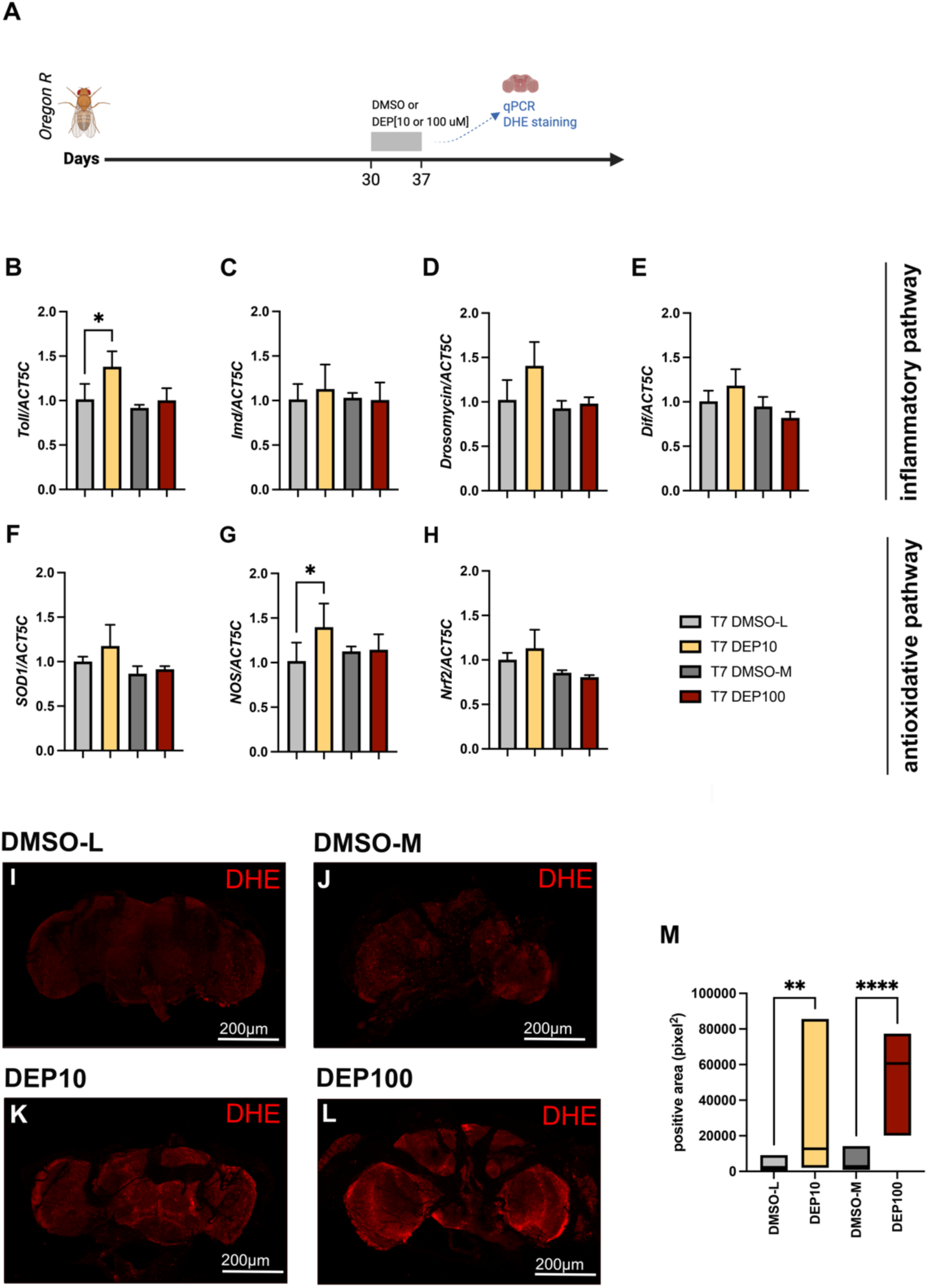
Effects of Diesel Exhaust Particles (DEPs) on aging flies following following acute exposure. (A) Schematic of acute DEP exposure (7 days) and analysis timeline. (B-H) Relative mRNA expression levels of neuroinflammatory and antioxidative markers in the brains of aging flies acutely exposed to DEP (DEP10and DEP100). Gene expression was normalized to *ACT5C*. The analyzed markers include (B) *Toll*, (C) *Imd*, (D) *Drosomycin*, (E) *Dif,* (F) *SOD1*, (G) *NOS*, (H) *Nrf2*. (I-L) Representative images of dihydroethidium (DHE) staining in fly brains indicating oxidative stress levels. (M) Quantification of DHE-positive areas in each condition. Statistical analysis was performed using the Kruskal-Wallis test followed by Dunn’s multiple comparison test (for comparisons among more than two groups). A p-value of < 0.05 was considered statistically significant *(*p < 0.05, **p < 0.01, ****p < 0.0001*). Each group included at least three biological replicates to ensure sufficient statistical power.

Next, we investigated the effects DEP on ROS production. We begun to meausre how aging itself affects the level of ROS in the brain. Flies were maintained in chemical-free condition until day 44 post eclosion and DHE staining was performed on D7, D14, D37 and D44 brains, respectively **(Supplementary Figure S4A).** As expected, the level of ROS gradually increases with aging, and the signal intensity was significantly noticed in the oldest group of flies (D44) **(Supplementary Figure S4A-F).** However, acute exposure to DEP either 10 or 100 markedly increased the ROS levels in treated flies in comparison to aged and vehicle-matched controls **(Figure 3I-M).** Moreover, such elevated ROS levels combined with locomotive functions alterations in old flies treated with DEP100 **(Supplementary Figure S2 J).** Moreover, similar to young flies following acute DEP exposure, aged flies also exhibited elevated ROS levels accompanied by altered locomotor function. However, unlike the hyperactivity observed in young flies, aged flies showed a decline in climbing ability **(Supplementary Figure S2 G).**

Overall, acute DEP exposure in aged flies led to a marked increase in brain ROS levels and severe locomotor impairments. These findings suggest that the brain responds to short-term DEP exposure in an age-specific manner, with aged flies being more vulnerable to oxidative stress induced by high DEP concentrations.

### DEP exposure alters BBB permeability and leads to long lasting brain damages in old flies

Since DEP exposure might link to the changing of BBB dynamics (37), we next investigated BBB integrity in aged flies following prolonged DEP exposure. We injected a high-molecular-weight dextran-conjugated fluorescent dye into the bodies of aged flies and then examined fluorescence in their brains **(Figure 4A)**. Vehicle control flies unexposed to DEP showed a very mild fluorescence, while DEP-exposed flies exhibited robust fluorescence signaling, indicating significantly compromised BBB integrity at both DEP10 and DEP100 exposure levels **(Figure 4B-F).**

**Figure 4.**
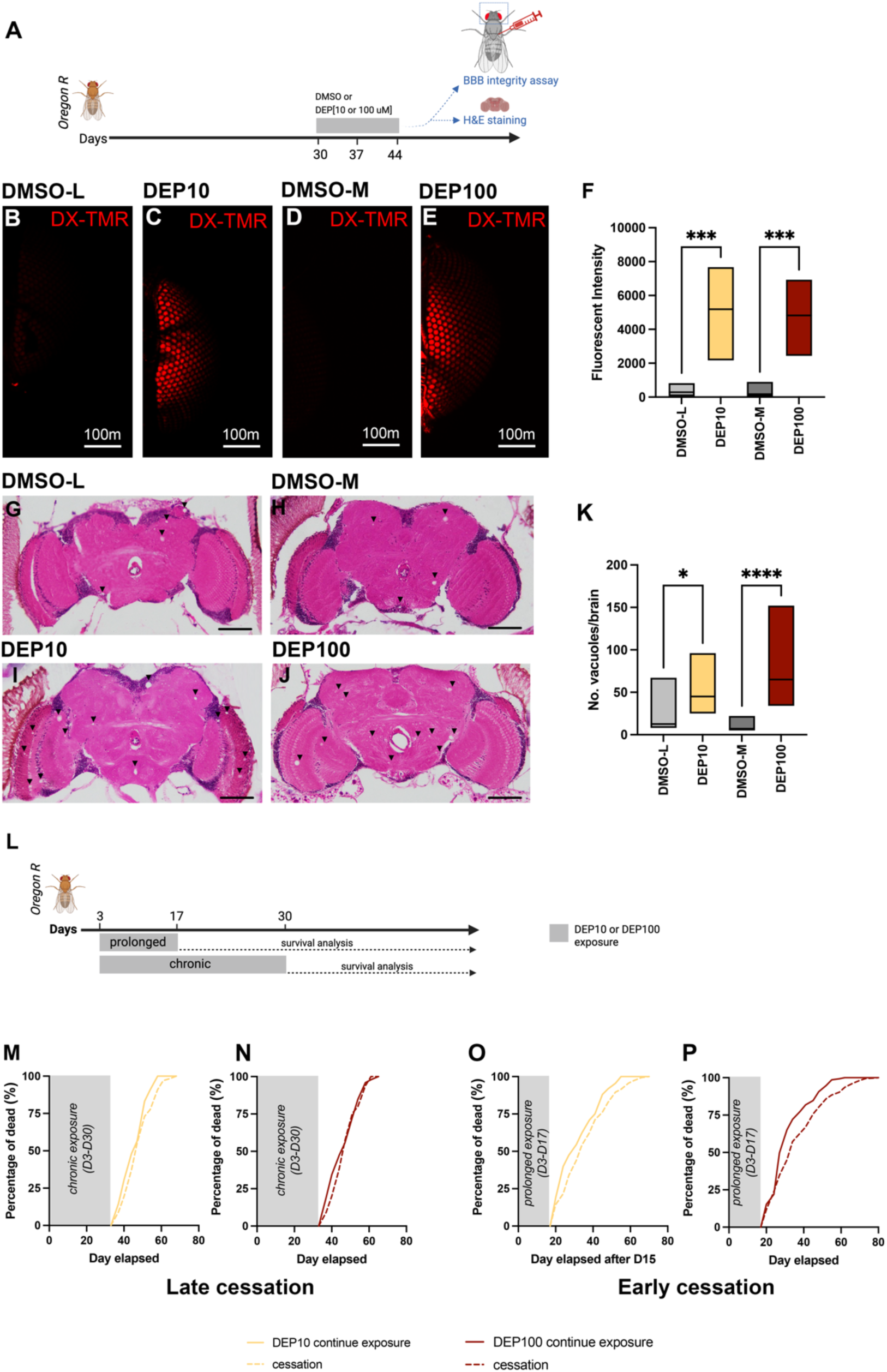
The detrimental effects of DEP exposure on brain structure and fly survival with or without cessation. (A) Schematic of prolonged DEP exposure and analysis timeline. (B-E) Blood-brain barrier (BBB) integrity in 30-day-old flies exposed to DEP10 (C) and DEP100 (E) for 14 consecutive days, compared to their respective vehicle controls, DMSO-L (B) and DMSO-M (D). BBB integrity was assessed using TMR (red) signals conjugated to 10,000 kDa Dextran. (F) Quantification of fluorescent intensity to compare BBB permeability in DEP10 vs DMSO-L and DEP100 vs. DMSO-M. (G-K) Representative images of hematoxylin and eosin (H&E)-stained brain sections from aging flies treated either with DMSO-L (G), DMSO-M (H), DEP10 (I), and DEP100 (J). Arrowheads indicate vacuoles, and scale bars represent 100 µm. (K) Quantification of vacuole formation in fly brains, expressed as the average number of vacuoles per brain in DEP10 vs. DMSO-L and DEP100 vs. DMSO-M. (A) Schematic of the experimental timeline for prolonged DEP exposure and cessation. (B–E) Percentage of deceased flies following chronic (B, D) or prolonged (C, E) exposure to DEP10 or DEP100, with or without a cessation period. Each group included at least 200 flies. Dashed lines indicate cessation; solid lines indicate continuous exposure without cessation. Data are presented as mean ± SD. Statistical analysis was performed using the Kruskal-Wallis test followed by Dunn’s multiple comparison test (for comparisons among more than two groups). Each group included at least ten adult brains to ensure robust statistical analysis. A p-value of < 0.05 was considered statistically significant *(*p < 0.05, **p < 0.01, ****p < 0.0001*).

To determine whether this BBB leakage was accompanied by permanent brain damage, we next examined brain structure. A significant increase in vacuolization was observed in old flies following prolonged DEP exposure at both DEP10 and DEP100 **(Figure 4I–K).** Although aged flies are intrinsically more prone to brain damage, as indicated by a small number of vacuoles after DMSO exposure **(Figure 4G-H, K)**, the extent of vacuolization was remarkably higher following DEP exposure **(Figure 4I–K).**

These data indicate that DEP exposure exacerbates age-dependent brain changes and suggest that the neuroinflammation and ROS accumulation observed in the brains of aged flies after prolonged DEP exposure may result from disruption of BBB integrity.

### DEP cessation is more beneficial when implemented at a young age

Chronic exposure to DEP is known to be detrimental to organisms (38, 39). In fact, in our experimental conditions, newly eclosed flies exposed to DEP throughout their lifespan exhibited significantly reduced survival, regardless of DEP concentration (**Supplementary Figure S5A–C).** Similarly, prolonged DEP exposure in aged flies also resulted in reduced lifespan, confirming the detrimental effects of long-term exposure across age groups **(Supplementary Figure S5D–E).**

To assess the effects of DEP exposure cessation, flies were initially subjected to either prolonged (14-day) or chronic (over 15-day) DEP exposure followed by a cessation period, and the percentage of dead flies was measured and compared to those continuously exposed without cessation **(Figure 4L).** Cessation after prolonged DEP exposure clearly improved fly survival, as the percentage of dead flies significantly decreased regardless of DEP concentration **(Figure 4O-P).** However, cessation following chronic exposure was less effective, particularly at higher concentrations **(Figure 4M,N).** These findings prompted us to investigate whether the age at the time of cessation influences its effectiveness. To explore this, we measured inflammatory pathway activation, antioxidant gene expression and ROS levels in the brains of flies exposed to acute DEP treatment followed by a 7-day cessation period, initiated at either 10 or 37 days of age **(Figure 5A, 7A).** These molecular markers, along with behavioral outcomes, were then compared to age-matched controls that did not undergo cessation **(Figure 5A, 7A).**

**Figure 5.**
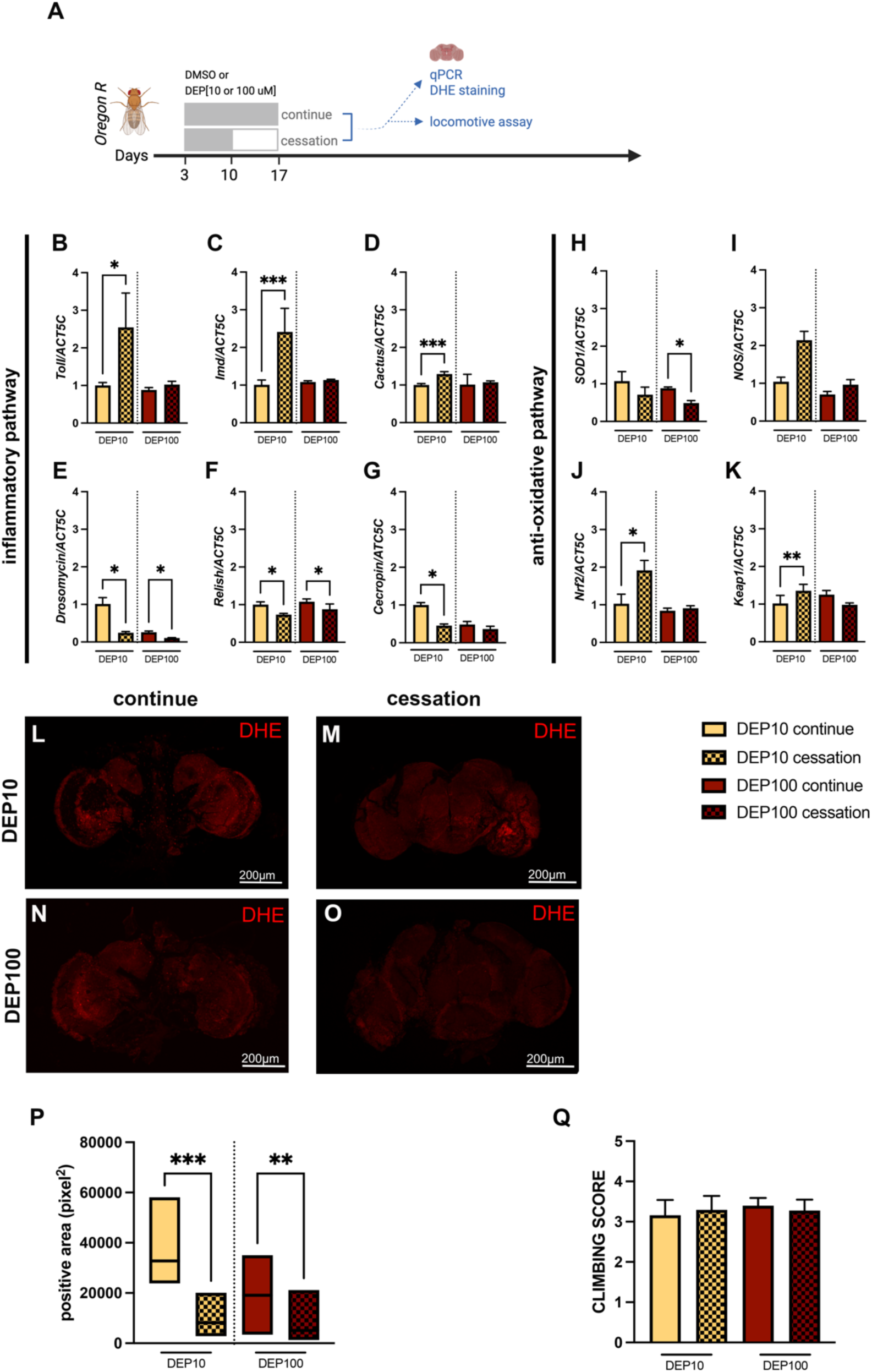
Effects of the cessation on young flies after DEP exposure. (A) Schematic of the experimental timeline for prolonged continue DEP exposure and cessation. Relative mRNA expression levels of neuroinflammatory (B-G) and antioxidative markers (H-K) in the brains of young adult flies prolonged exposed to DEP10 or DEP100 or had 7-day cessation period after 7-day exposure (cessation). Markers analyzed and normalized to *ACT5C* include (B) *Toll*, (C) *Imd*, (D) Cactus, (E) *Drosomycin*, (F) *Relish (G) Cecropin*, (H) *SOD1*, (I) *NOS*, and (J) *Nrf2* and (K) *Keap1.* (L-O) Representative images showing dihydroethidium (DHE) staining in fly brains with either prolonged exposure or underwent to 7-day cessation after the exposure indicating oxidative stress levels. (P) Quantification of DHE-positive area in each condition. Locomotor performance following DEP exposure regimen (Q) Statistical analysis was performed using the Kruskal-Wallis test followed by Dunn’s multiple comparison test (for comparisons among more than two groups). A p-value of < 0.05 was considered statistically significant *(*p < 0.05, **p < 0.01, ***p < 0.001*). Each group included at least three biological replicates to ensure sufficient statistical power.

Interestingly, cessation of DEP exposure in young flies led to a reduction in the expression of key downstream effectors of inflammatory pathways, including *Drs*, *Relish*, and *Cecropin*, along with their inhibitor *Cactus*, indicating a potential attenuation of neuroinflammation. This effect was evident in both DEP10– and DEP100-treated groups compared to controls **(Figure 5B–G).** A compensatory antioxidant response could be also followed by a cessation of DEP10, as upregulation of the master regulators of antioxidative defense like *Nrf2* and *Keap1* was observed in comparison to no cessation group **(Figure 5J–K).** However, no changes were observed in *SOD1 and NOS* **(Figure 5H-I).** In contrast, cessation from DEP100 appeared to have a less beneficial effect on the antioxidative response, with no observable changes in most markers and a notable reduction in *SOD1* expression **(Figure 5H).** Moreover, cessation in young flies exposed to either DEP10 or DEP100 also significantly reduced brain ROS levels **(Figure 5L–P),** despite no major changes being observed in locomotor function **(Figure 5Q).**

In contrast to young flies, cessation from DEP exposure in aged flies **(Figure 6A)** was less effective in reducing neuroinflammation. This was evidenced by the continued upregulation of *Drosomycin* and the absence of changes in *Relish* or *Cecropin*, consistent with sustained expression of the upstream regulators *Toll* and *Imd* **(Figure 6B–G).** Furthermore, the downregulation or lack of change in *Nrf2* and *Keap1* suggests that cessation in aged flies may fail to activate an effective antioxidative defense response **(Figure 6H–K).** Finally, a reduction in brain ROS levels was also observed in aged flies following cessation from either DEP10 or DEP100 exposure **(Figure 6L–P),** although no significant changes were detected in locomotor function **(Figure 6Q).**

**Figure 6.**
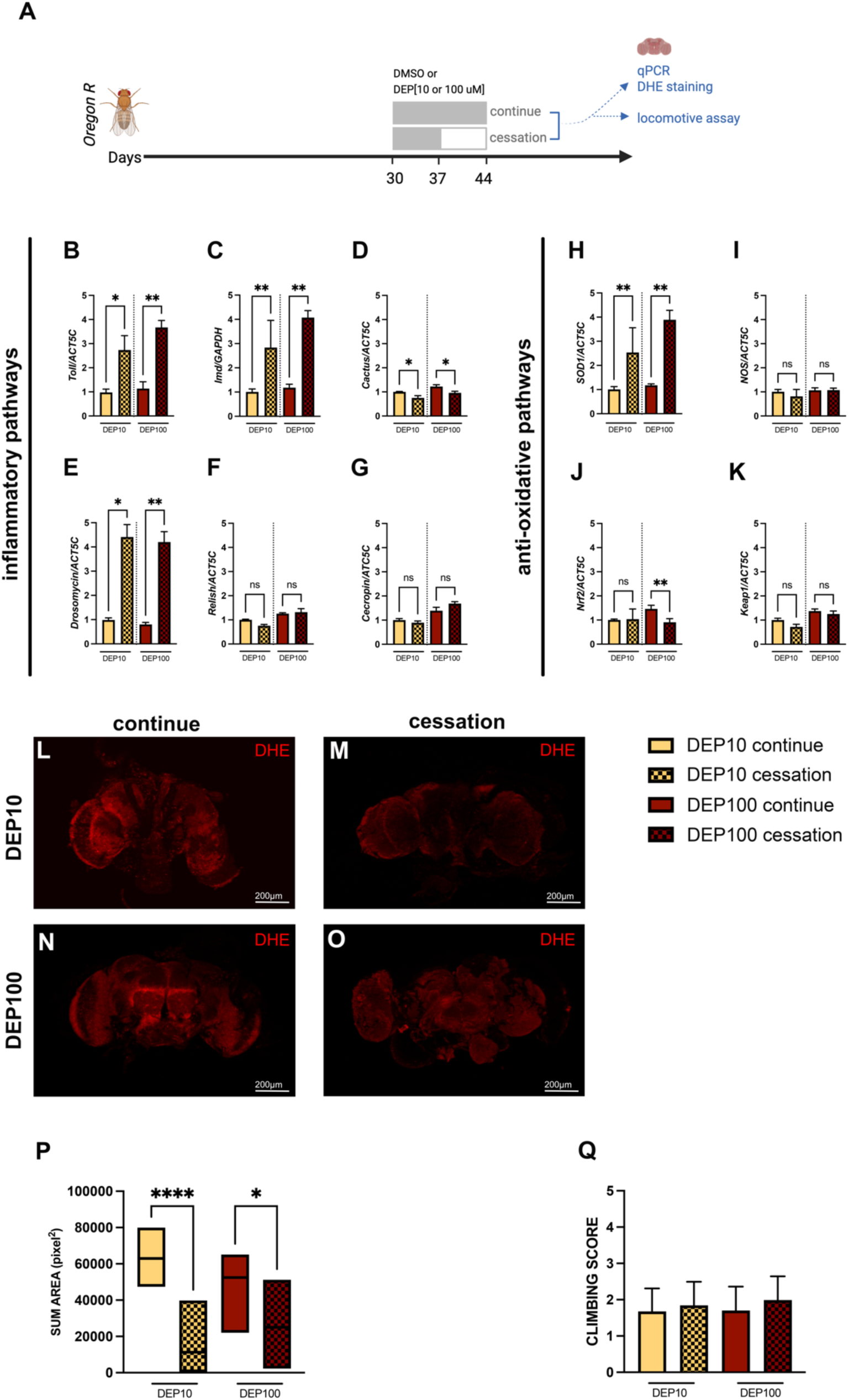
Effects of the cessations on on aging flies after DEP exposure. (A) Schematic of the experimental timeline for prolonged continue DEP exposure and cessation. Relative mRNA expression levels of neuroinflammatory (B-G) and antioxidative markers (H-K) in the brains of young adult flies prolonged exposed to DEP10 or DEP100 or had 7-day cessation period after 7-day exposure (cessation). Markers analyzed and normalized to *ACT5C* include (B) *Toll*, (C) *Imd*, (D) Cactus, (E) *Drosomycin*, (F) *Relish (G) Cecropin*, (H) *SOD1*, (I) *NOS*, and (J) *Nrf2* and (K) *Keap1*. (L-O) Representative images showing dihydroethidium (DHE) staining in fly brains with either prolonged exposure or underwent to 7-day cessation after the exposure indicating oxidative stress levels. (P) Quantification of DHE-positive area in each condition. Locomotor performance following DEP exposure regimen (Q) Statistical analysis was performed using the Kruskal-Wallis test followed by Dunn’s multiple comparison test (for comparisons among more than two groups). A p-value of < 0.05 was considered statistically significant *(*p < 0.05, **p < 0.01, ***p < 0.001*). Each group included at least three biological replicates to ensure sufficient statistical power.

Overall, cessation from DEP exposure is beneficial in attenuating neuroinflammation and enhancing antioxidant defenses when implemented at a young age. However, these protective effects are markedly reduced when cessation occurs in older flies. Nonetheless, cessation consistently reduces brain ROS levels, regardless of the age at the time of cessation.

## Discussion

Our study reveals that even brief, acute exposure to diesel exhaust particles (DEP), a major component of urban air pollution can induce lasting, age-dependent neurotoxicity. While earlier research has linked environmental pollutants to general brain vulnerability, our data show that aged brains are particularly susceptible, exhibiting persistent oxidative stress, impaired blood-brain barrier (BBB) integrity, and reduced recovery following exposure. In contrast, younger brains demonstrate greater resilience and partial reversal of DEP-induced effects.

We found that acute DEP exposure significantly altered brain gene expression, including early activation of inflammatory and antioxidant pathways, even at low doses and without visible toxicity. These molecular changes were accompanied by increased reactive oxygen species (ROS) and locomotor impairments in young flies, suggesting that even limited exposure can initiate functional and oxidative stress-related deficits prior to overt neurodegeneration. ROS, therefore, may serve as an early biomarker for pollutant-induced neurotoxicity.

*Drosophila* is known to be highly sensitive to environmental toxins such as metals and organic pollutants (40), and our study confirms that DEP elicits strong physiological responses in this model. Prior studies have shown that airborne pollutants can disrupt genetic stability, inflammatory signaling, and metabolism in flies, regardless of exposure route whether via inhalation, ingestion, or injection (41–45). Despite using an oral administration route, which involves systemic processing before reaching the brain, we observed strong neurotoxic outcomes. This suggests that DEP may act either directly on neural tissue or indirectly via systemic inflammatory and oxidative pathways. Additionally, oral DEP exposure has been shown to disrupt gut microbiota, potentially heightening immune activation via Imd-dependent pathways and antimicrobial peptide expression (46, 47).

A key finding of our study is that age significantly shapes the brain’s response to DEP exposure. Aged flies exhibited a more severe phenotype, including a distinct transcriptional profile, greater ROS accumulation, and marked behavioral decline. These results suggest a diminished capacity in the aging brain to counteract environmental stressors. We also observed increased BBB permeability and vascular-like brain alterations in aged flies, suggesting a direct route for DEPs and toxins into the brain, exacerbating microglial activation and neuroinflammation. This mechanism is consistent with known links between pollution, oxidative stress, and neurodegenerative pathologies (48–50).

Finally, while cessation of DEP exposure benefited both age groups, only young flies showed significant improvements in molecular markers and behavioral performance. In aged flies, cessation failed to reverse most neurotoxic effects, likely due to cumulative or irreversible brain damage. These findings underscore a critical window of neuroplasticity in early life, which narrows with age, and highlight the importance of minimizing pollutant exposure early in life to preserve long-term brain health.

## Conclusions

In conclusion, our findings demonstrate that acute DEP exposure compromises brain health in an age-dependent manner, causing oxidative stress, behavioral deficits, and lasting structural damage, particularly in aged individuals. This study reinforces the role of environmental pollutants as risk factors for age-related brain diseases and emphasizes the importance of early-life intervention. *Drosophila* offers a powerful model to dissect the molecular underpinnings of pollution-induced neurotoxicity and to identify age– and stage-specific protective mechanisms. Future studies should aim to elucidate the long-term trajectory of these changes, evaluate chronic low-dose exposures, and explore potential therapeutic or preventive strategies.

## CRediT authorship contribution statement

Conceptualization: S.J. and L.LP; Data curation: S.J. and R.Y.; Formal Analysis: S.J, R.Y.and L.LP.; Funding acquisition; S.J. and R.Y.; Investigation: S.J., R.Y., S.P., T.Y., K.P. and P.M.; Methodology: S.J. and L.LP; Project administration: S.J.; Resources; S.J; Software: R.J. and L.LP.; Supervision; L.LP and S.J.; Validation: S.J., and L.LP.; Visualization: R.Y. and S.J.; Writing – original draft: S.J. Y.N. and L.LP.; Writing – review & editing: All authors have reviewed and approved the final version of this manuscript.

## Acknowledgements

We would like to express our gratitude to the technical staffs of Pharmacology department, Anatomy Department and Center of Multidisciplinary Technology for Advanced Medicine (CMUTEAM), Faculty of Medicine, Chiang Mai University, for their generous support. In particular, we acknowledge Miss Thunpitcha Meesawat for administrative assistance and sincerely thank Mr. Papon Hitmool, Mr. Sakorn Phongchankhiao, Ms Ratchanaree Suepsaipaeng, Ms Chansunee Panto and Mr. Tawan Munwanna for technical assistance. We are also thank the Bloomington Stock Center for providing fly strains, the Science and Educational Company Limited (SCIED) for training in confocal scanning electron microscopy.

## Data Availability

The authors confirm that the data supporting the findings of this study are available within the article and its supplementary material.

