## supplementary figures for "Diesel exhaust particles induce lasting and age-dependent damage to the brain in *Drosophila melanogaster*"

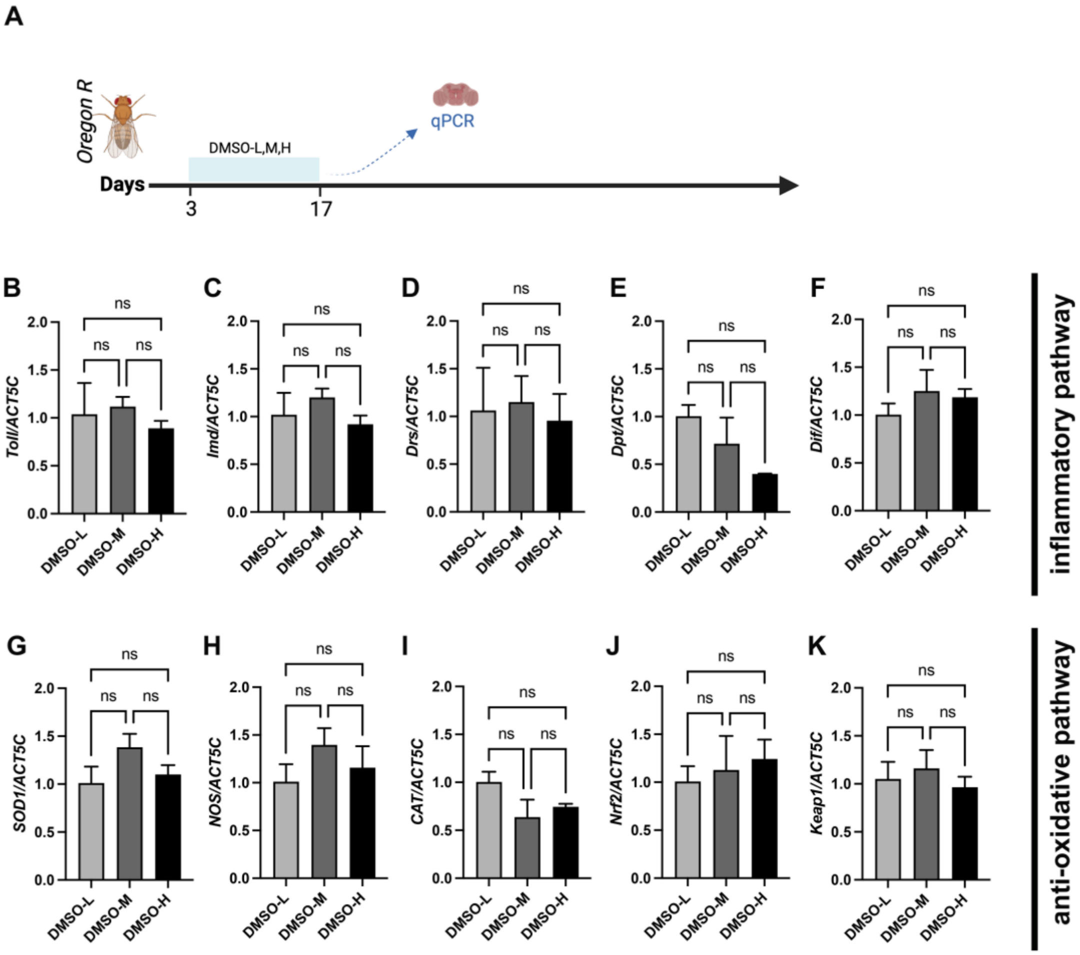


**Supplementary Figure 1 mRNA expression profiles in the heads of young flies exposed to varying concentrations of DMSO.**

(A) Schematic representation of the 14-day DMSO exposure in young flies. (B–K) Relative mRNA expression of genes involved in inflammatory pathways: (B) Toll, (C) *Imd*, (D) *Drosomycin (Drs)*, (E) *Dpt (Dipterician)*, (F) *Dif*, and anti-oxidative pathways: (G) *SOD1*, (H) *NOS*, (I) *CAT*, (J) *Nrf2*, (K) *Keap1*. Flies were exposed to DMSO at three concentrations: DMSO-L (0.05%), DMSO-M (0.5%), and DMSO-H (1%). mRNA levels were normalized to *ACT5C* and expressed as mean ± SD. Statistical analysis was performed using the Kruskal–Wallis test followed by Dunn's multiple comparisons. A p-value $< 0.05$ was considered statistically significant (ns = not significant). Each group included at least three biological replicates.

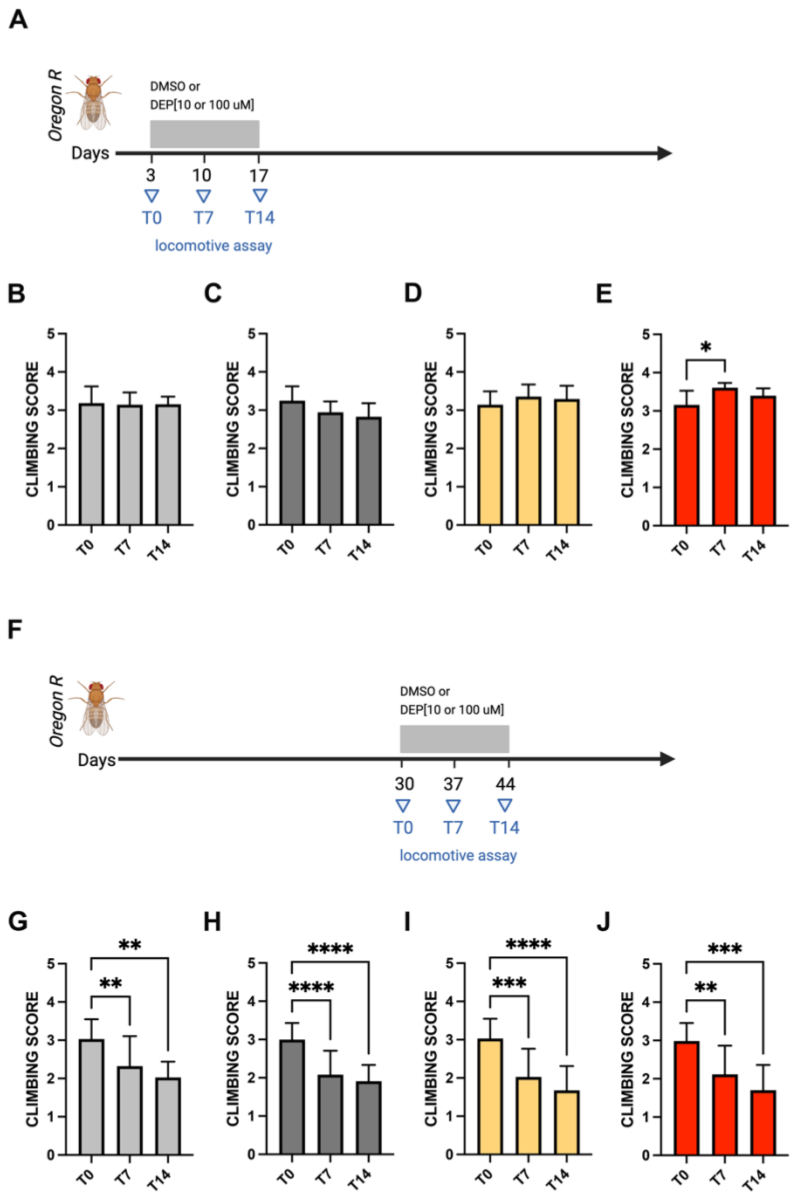


**Supplementary Figure 2 Locomotor ability of young and aging flies exposed to DEPs or DMSO.** (A) Schematic representation of young flies treated with DMSO-L, DMSO-M, DEP10, or DEP100, assessed at baseline (T0), after 7 days (T7), and after 14 consecutive days (T14) of exposure. (B–E) Climbing scores of young flies treated with DMSO-L (B), DMSO-M (C), DEP10 (D), and DEP100 (E). (F) Schematic representation of aging flies treated with DMSO-L, DMSO-M, DEP10, or DEP100, assessed at T0, T7, and T14. (G–J) Climbing scores of aging flies treated with DMSO-L (G), DMSO-M (H), DEP10 (I), and DEP100 (J). Climbing scores are presented as mean ± SD. Statistical analysis was performed using the Kruskal–Wallis test followed by Dunn's multiple comparison test (for comparisons among more than two groups). A p-value of < 0.05 was considered statistically significant (*$*p < 0.05$, $**p < 0.01$, $***p < 0.001$, $****p < 0.0001$*, ns = not significant). Each group included at least three biological replicates, with a total of 150 flies per treatment group to ensure sufficient statistical power.

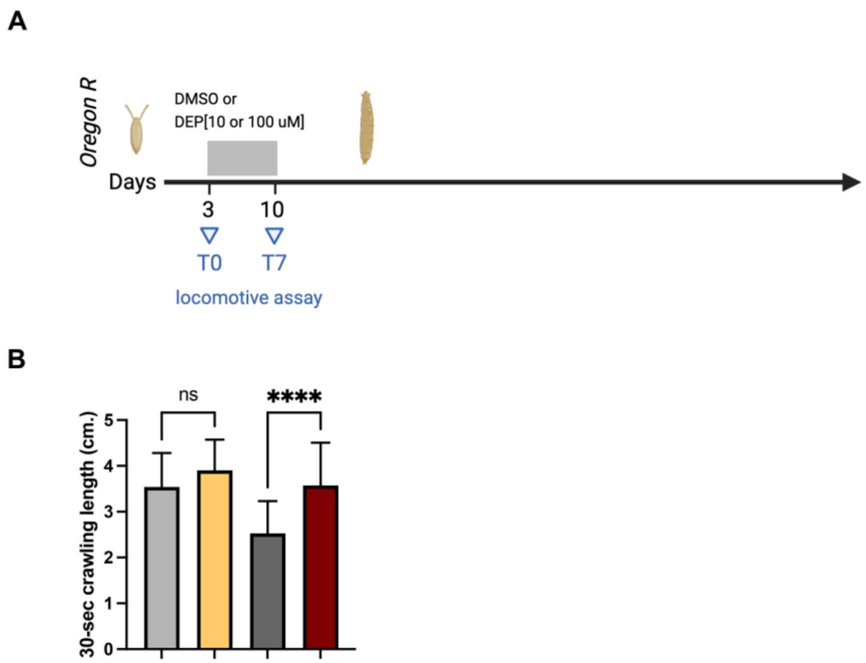


**Supplementary Figure 3. Locomotor Ability of Third Instar Larvae**

(A) Schematic representation of third instar larvae treated with DMSO-L, DMSO-M, DEP10, or DEP100 for 7 days, starting from the embryonic stage. (B) Crawling distance (cm) of larvae after 7-day exposure. Crawling distances are presented as mean ± SD. Statistical analysis was performed using the Kruskal–Wallis test followed by Dunn's multiple comparison test (for comparisons among more than two groups). A p-value < 0.05 was considered statistically significant (*$****p < 0.0001$*). Each group included at least three biological replicates, with a total of 50 larvae per treatment group to ensure sufficient statistical power.

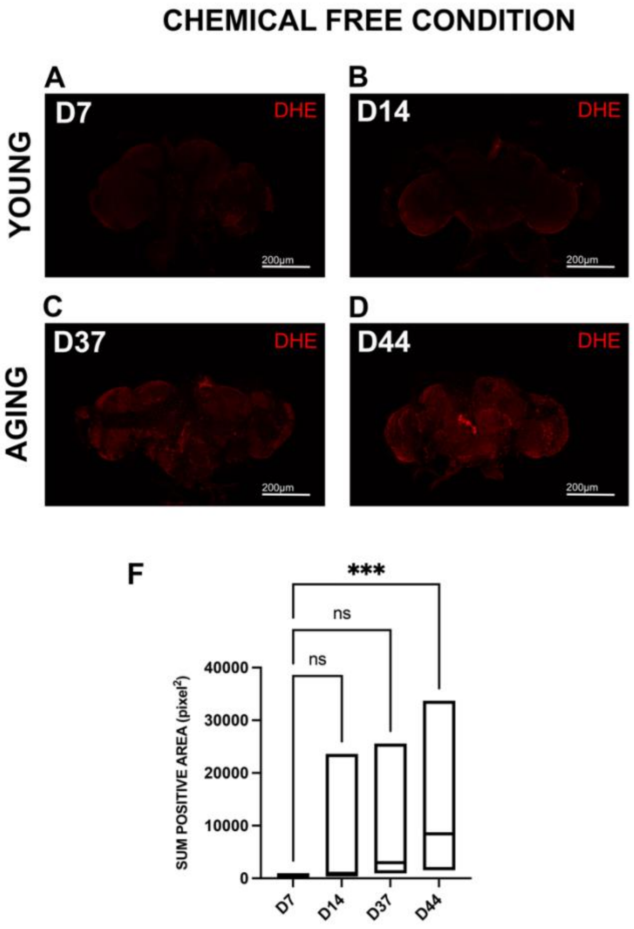


**Supplementary Figure 4 Age-Related Baseline ROS Levels in Fly Brains**

(A–D) Representative images of dihydroethidium (DHE) staining in fly brains at different ages, indicating oxidative stress levels: (A) 7-day-old, (B) 14-day-old, (C) 37-day-old, and (D) 44-day-old flies. (E) Quantification of DHE-positive areas across different age groups. Data are presented as mean ± SD and displayed in a box plot. Statistical analysis was performed using the Kruskal–Wallis test followed by Dunn's multiple comparison test. A p-value < 0.05 was considered statistically significant *(***p < 0.001*; ns = not significant). Each group included at least three biological replicates to ensure sufficient statistical power.

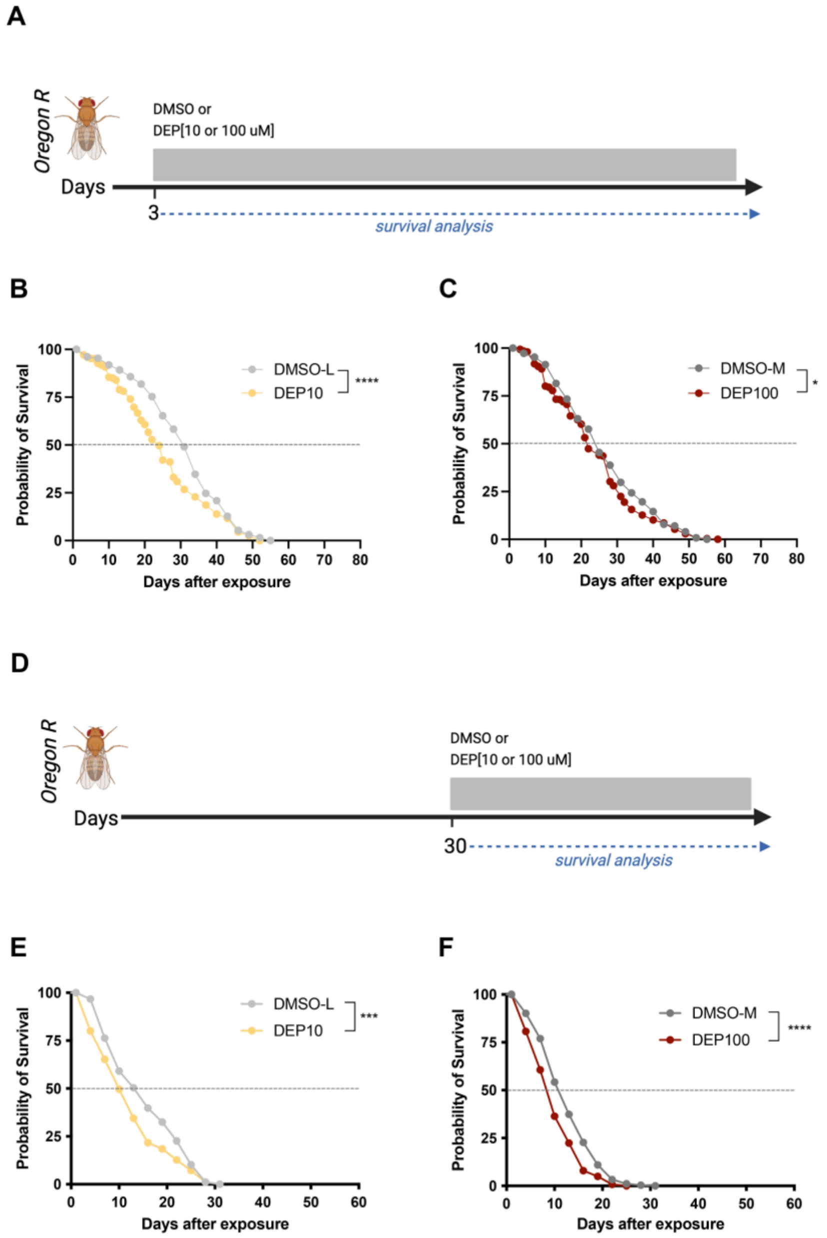


**Supplementary Figure 5 Lifespan Analysis of Young and Aging Flies Exposed to either DEPs or DMSO** (A) Schematic representation of young flies exposed to DEP10, DEP100 or their respective DMSO control. Lifespan of young flies exposed to (B) DEP10 or (C) DEP100 compared to their DMSO controls (D) Schematic representation of aging flies exposed to DEP10, DEP100 or their respective DMSO control. Lifespan of young flies exposed to (E) DEP10 or (F) DEP100 compared to their DMSO controls. Survival curves validated by Log-rank (Mantel-Cox) test. A p-value of < 0.05 was considered statistically significant $**p < 0.01$, $***p < 0.01$, $****p < 0.0001$. Each group included a total of 200 flies per treatment group to ensure sufficient statistical power.

**Supplementary Table 1 List of Primers used for qRT-PCR.**

| Gene Name | Forward Sequences (5'-3') | Reverse Sequences (5'-3') |
|---|---|---|
| *Toll* | *GTTCAGATGCGACGGTGTATG* | *GCCGTGTTATGTTCATGCCC* |
| *Imd* | *AGGGACGCCTGGAAAAGGA* | *GGATTCGGTCAGTCCGAGGA* |
| *Drosomycin* | *CTGGGACAACGAGACCTGTC* | *ATCCTTCGCACCAGCACTTC* |
| *Diff* | *CAGTGTGGCAGGAGCTGTT* | *GGAATGTGCGACGGCATTAC* |
| *dSOD1* | *GGACCGCACTTCAATCCGTA* | *TGGAGTCGGTGATGTTGACC* |
| *NOS* | *ATGTCGCAGCATTTCACATCG* | *GGCGTTGCTTGAGTTTTGATTT* |
| *Nrf2* | *TTACATCTACGAGTACGCCGC* | *ACTGGAGCTCAAAACCGCTAA* |
| *Keap1* | *CCACCGTGGAGCGTTATGATA* | *TTCCTGCATTCTGGACCAAGG* |
| *Cactus* | *AACCACTGATTCGGGCTTCAT* | *CCTGCTGATCCTTATCCTGTTCC* |
| *Relish* | *GGTGATAGTGCCCTGCATGT* | *CCATACCCAGCAAAGGTCGT* |
| *Cecropin* | *AAGCCGGTTGGTGAAGAAA* | *GTCCTTGAATGGTTGCATCCC* |
| *ACT5C* | *CAACTGGGACGATATGGAGAAG* | *GTCTCGAACATGATCTGGGTC* |
